# Mechanical characterisation of the developing cell wall layers of tension wood fibres by Atomic Force Microscopy

**DOI:** 10.1101/2021.09.23.461481

**Authors:** Olivier Arnould, Marie Capron, Michel Ramonda, Françoise Laurans, Tancrède Alméras, Gilles Pilate, Bruno Clair

## Abstract

Trees generate mechanical stresses at periphery of stem and branches to improve their strength and to control the orientation of their axes. This key factor in the biomechanical design of trees, named “maturation stress”, occurs in wood fibres during cellular maturation when their secondary cell wall thickens. In this study, the spatial and temporal stiffening kinetics of the different cell wall layers were recorded during fibre maturation on a sample of poplar tension wood using atomic force microscopy. The thickening of the different layers was also recorded. The stiffening of the CML, S_1_ and S_2_-layers was initially synchronous with the thickening of the S2-layer and continued a little after the S2-layer reached its final thickness as the G-layer began to develop. In contrast, the global stiffness of the G-layer, which initially increased with its thickening, was close to stable long before it reached its final maximum thickness. A limited radial gradient of stiffness was observed in the G-layer, but it decreased sharply on the lumen side, where the new sub-layers are deposited during cell wall thickening. Although very similar at the ultrastructural and biochemical levels, the stiffening kinetics of the poplar G-layer appears to be very different from that described in maturing bast fibres.

## Introduction

Wood fibres have mechanical functions in the living tree. Mature wood fibres give the tree axis sufficient stiffness and strength to withstand its own weight and additional loads such as wind, snow/ice or fruit (Niklas, 1992). In addition to this “skeletal” function, wood fibres also have a “muscular” function that has two major goals. First, it allows the tree to control its posture by actively generating forces that can bend the axes (stem, branches) upwards or compensate for the effect of gravity (Alméras and Fournier, 2009; Fournier *et al*., 2014; Moulia *et al*., 2006; Scurfield, 1973). Second, it improves axes resistance to bending loads, such as wind, by a beneficial stress profile *(i.e*., tensile longitudinal stress at the periphery of the stem) (Bonser and Ennos, 1998; Alméras *et al*., 2018). These mechanical stresses originate from physico-chemical changes of the fibre cell wall that would result in major deformation if they were not prevented by the older, stiff tissue, surrounding them. In place of strain, this leads to the development of a high mechanical stress named “maturation stress” or “growth stress” (Archer, 1986). These fibre properties are progressively built up during their development.

The development of wood fibres is usually described in three phases: division taking place in the cambium, extension during which each cells reached its final size and shape, and maturation during which the secondary wall is developing. During this last maturation phase, constitutive polymers are progressively added to the wall from the cytoplasm, leading to the secondary wall thickening. The kinetics of fibre extension and cell wall thickening are well known and can be deduced from observations on transverse section using optical microscopy (Abedini *et al*., 2015; Andrianantenaina *et al*., 2019; Pérez-de-Lis *et al*., 2022). However, the initial mechanical state of the deposited polymers, the evolution of their mechanical properties or their stress state during maturation are more difficult to measure and almost no data are available in the literature on these parameters. Data collection of time and space changes in the mechanical properties of the secondary wall is a first step toward a better understanding on the links between wood maturation and the building of wood quality. Here, we propose to measure these changes in the special case of tension wood formation.

Tension wood is produced by angiosperms and characterised by a strong tensile stress in the fibre axis direction of the order of several tens of MPa. In tension wood, mechanical stress is known to be mainly generated within a specific gelatinous cell wall layer, named the G-layer (Côté *et al*., 1969; Dadswell and Wardrop, 1955; Fang *et al*., 2008; Ghislain and Clair, 2017; Onaka, 1949). Fibres containing a gelatinous layer are widespread in the plant kingdom and can be present in various organs and tissues (Gorshkova *et al*., 2018). Depending on their cell wall composition, fibres can be classified either as lignin-rich (*i.e*., wood fibres) or virtually lignin-free, such as part of tension wood fibres (Ghislain *et al*., 2019) or bast fibres found in flax, hemp, ramie, nettle, etc (Gorshkova and Morvan, 2006). It is this latter category of fibres that is often named gelatinous fibre. Some authors have proposed to call this gelatinous layer a tertiary cell wall but a dedicated article exposes several arguments arguing that this term is not appropriate (Clair *et al*., 2018). While poplar tension wood fibres are of secondary origin, flax fibres are primary phloem fibres, or bast fibres. However, these gelatinous fibres have several similarities in the biochemical composition of their cell walls with a high content of crystalline cellulose oriented parallel to the fibre axis, very little or no lignin as already mentioned, a matrix of non-cellulosic polysaccharides rich in pectic ß-(1-4)-galactans and, to a lesser extent, of type II arabinogalactan. The same is observed when comparing the transcriptome of developing tension wood fibres and of flax phloem fibres, with multiple, distinct chitinases, β-galactosidases, arabinogalactan proteins and lipid transfer proteins. Compare for example Roach *et al*. (2007) and Déjardin *et al*. (2004). Moreover, the thickening of the cell wall of flax fibres involves considerable remodelling of the deposited layers: indeed, the newly formed layers of the secondary cell wall of developing flax fibres, referred to as the immature Gn-layer, have a loose structure that will get more and more compact and stiff during their maturation toward a G-layer (Goudenhooft *et al*., 2018; Petrova *et al*., 2021).

The mechanisms responsible for the generation of high tensile stress during G-layer maturation are still the subject of debate. Several hypothetical models have been proposed, which are reviewed in Alméras and Clair (2016). Gaining knowledge on the chemical, physical and mechanical states of the constitutive materials and their changes during cell wall maturation have proven particularly useful in distinguishing between these models. For example, it has been observed that the G-layer contains mesopores of several nanometres (Chang *et al*., 2009; Clair *et al*., 2008), and that these pores swell during maturation (Chang *et al*., 2015). It has also been shown that crystalline microfibrils are progressively put under tension during maturation (Clair *et al*.,2011). The synchronicity between these two phenomena supports the hypothesis that pore swelling is related to the induction of tensile stress in the crystalline microfibrils and thus to maturation stresses in the G-layer (Alméras and Clair, 2016). A crucial factor is the change in cell wall stiffness during maturation. Indeed, using mechanical modelling, it has been shown that the relative kinetics of stiffening and stress induction affect the resulting state of stress in the tree (Alméras *et al*., 2005; Pot *et al*., 2014; Thibaut *et al*., 2001). As reported by Thibaut *et al*. (2001), the tendency of the material to deform in response to physico-chemical changes can result in stress of high magnitude only if the cell wall is already sufficiently stiff. To the best of our knowledge, information on the stiffening kinetics of wood cell wall layers is currently lacking and the only measurements available are at the tissue scale (Grozdits and Ifju, 1969; Pot *et al*., 2013a; 2013b).

One of the most promising and frequently used techniques today, nanoindentation, probes the mechanical properties at the cell wall scale. It enables access to the mechanical properties within the cell wall layers with modifications reduced to a minimum. This technique has already been used to estimate the indentation modulus of mature native or thermo-mechanically modified cell walls of wood fibres (Eder *et al*., 2013), lignifying spruce tracheid secondary cell walls (Gindl *et al*., 2002) and (thick) fibre cell walls within a maturing vascular bundle of bamboo (Huang *et al*., 2016; Wang *et al*., 2012). However, as widely recognized in the case of metallic materials, the radius of the plastically affected volume around the indenter is about three times the residual indent size for an isotropic material, and even more for the elastically affected one (Johnson 1987; Sudharshan Phani and Oliver, 2019). This technique therefore requires a layer thickness at least three times the size of the indent, which are typically in the micrometre range, to avoid measurement artefacts (Jakes *et al*., 2009). As the width of the cell wall layers in the developing and maturation stages vary from almost zero (cambium, beginning of the layer deposition) to a few micrometres (mature S2 and/or G-layer), interpreting the measurements obtained by nanoindentation in the presence of a gradient of properties or within a thin layer is not straightforward, nor possible close to the cambium, due to boundary effects. In such cases, atomic force microscopy (AFM) appears to be the best way to perform mechanical measurements within each cell wall layer (Arnould and Arinero, 2015; Casdorff *et al*., 2017; 2018; Clair *et al*., 2003, Coste *et al*., 2021; Nair *et al*., 2010; Normand *et al*., 2021). This technique has already been used to investigate, for example, the development of bast fibres within a flax stem (Goudenhooft *et al*., 2018) and of the primary cell walls in the inner tissues of growing maize roots (Kozlova *et al*., 2019).

The aim of the present work was to measure changes in the indentation modulus of each cell wall layer during the maturation of poplar tension wood fibres using AFM. As it was not possible to monitor the maturation of a single cell over time, as a proxy, we chose to perform measurements on several cells in the same row, from cambium to mature wood, that were therefore at different stages of development. Using the kinetics of cell wall thickening as a basis for comparison, the stiffening of the different layers of the cell wall was compared to other known phenomena such as changes in mesoporosity and in crystalline cellulose strain. In addition, thanks to the nanometric spatial resolution of AFM measurements, we investigated G-layer stiffening during thickening, *i.e*., the kinetics of stiffening within the G-layer, and fluctuations in the mechanical states of a new freshly deposited sub-layer. Finally, the kinetics and stiffness gradient of the poplar G-layers were compared with data available in the literature on flax phloem fibres containing a gelatinous layer.

## Materials and methods

### Sample preparation

The experiments were conducted on a wood sample cut out of a young hybrid poplar tree (*Populus tremula × Populus alba*, INRA clone 717-1-B4) tilted to induce the production of tension wood. This clone was chosen as it is easy to multiply and has therefore already been used for several studies related to tension wood formation (Abedini *et al*., 2015; Guedes *et al*., 2017; Lafarguette *et al*., 2004). This hybrid poplar plant was grown upright in a controlled greenhouse (located at INRAE, Orléans, France) for two months before inducing the formation of tension wood on the upper side of its stem, by tilting the plant 30° from the vertical and holding it in this position by binding the stem to a rigid stick. No up-righting process was thus allowed, which ensured the formation of tension wood along almost all the length of the stem and during the whole tilting period. Twenty-two days after tilting, a 5-cm long stem section (estimated diameter 1 cm) was collected at the base of the stem, at around 10 cm above the ground. Small wood sub-samples, a few mm in size, were cut out of the tension wood side and fixed for 4 h in 2.5% formaldehyde and 0.1% glutaraldehyde in 0.1M McIlvaine citrate-phosphate buffer, pH 6.8, followed by 3 × 10 min under moderate vacuum. After thorough rinsing in McIlvain buffer, the sample was partially dehydrated in increasing series (25%, 50%, 70%) of ethanol and progressively impregnated with LR-White medium grade resin (Agar Scientific Ltd, Stansted, UK), in a series of resin and ethanol mixes containing a progressively increasing percentage of resin (20% 2 h, 40% 4 h, 60% 4 h, 80% 24 h, 100% 2+8 days). During the last pre-embedding step, in pure resin, the sample was placed under moderate vacuum for 3 × 10 minutes. Finally, the samples were embedded in gelatine capsules filled with pure resin and heated in an oven at 56°C for 24 h for polymerization. Semi-thin transverse sections (0.5 to 1 μm) were cut with a Histo diamond knife (Diatome Ltd, Nidau, Switzerland) installed on a Ultracut S microtome (Leica Microsystems, Vienna, Austria) to trim the block. To avoid the deformation commonly observed in G-layers as a result of their swelling, detachment and collapse after stress release (Clair *et al*., 2005a; 2005b), at least the first 50 μm of the sample were trimmed and discarded. Finally, very thin sections (about 50 nm thick in the last step) were made at a low cutting speed (≈1 mm/s) using an Ultra AFM diamond knife (Diatome Ltd, Nidau, Switzerland) to obtain a nearly perfect flat surface. AFM measurements were carried out on the remaining block.

### Optical measurement of the cell wall layer thickness

After the AFM experiments, semi-thin transverse sections (0.9 μm) were cut with a Histo diamond knife (Diatome Ltd, Nidau, Switzerland) installed on an Ultracut R microtome (Leica Microsystems SAS, Nanterre, France). These sections were stained using Richardson’s azur II and methylene blue (Richardson *et al*., 1960) and mounted on slides using Canada balsam. The slides were observed under a light microscope (DMLP, Leica Microsystems SAS, Nanterre, France) with immersion oil lenses (Fig. 1). Richardson’s staining makes possible an easy detection of the presence of the G-layer in wood fibres of poplar, proof of the occurrence of tension wood. Conversely, the absence of G-layer indicates that no tension wood was present, as can be observed on the wood formed before tilting (on the right side in Fig.1). This has been confirmed by the absence of an additional G-layer in the following AFM observations.

**Figure 1:**
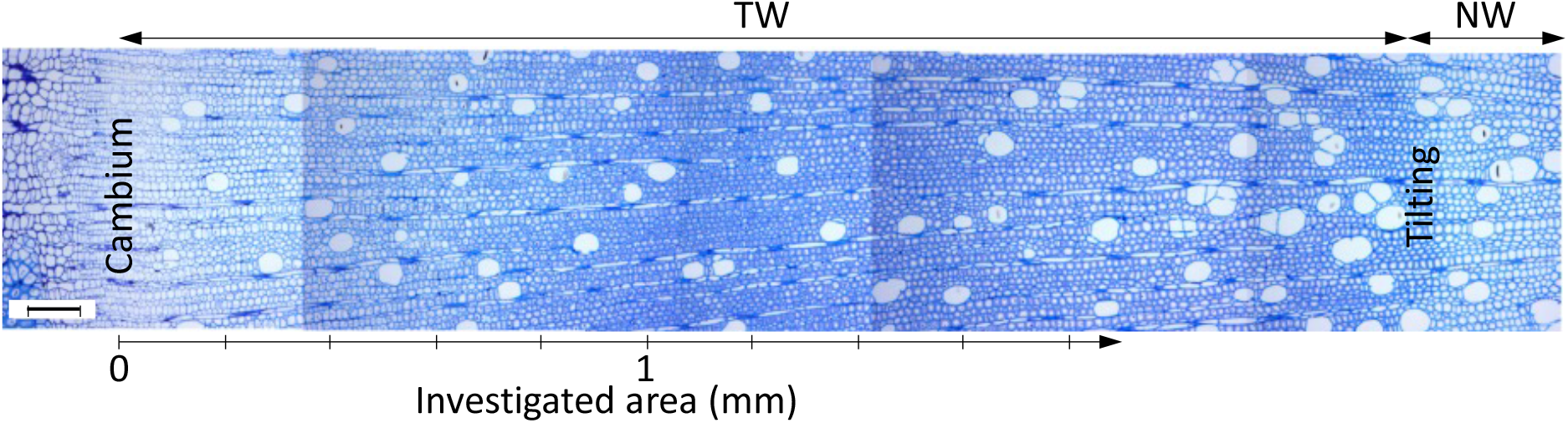
optical image of the transverse section of the wood sample (Richardson’s staining) with the tension wood (TW) area between the cambium and the normal wood (NW) produced before the tree was tilted. The reference distance from the cambium was measured approximately in the middle of the cambial zone. Vertical lines correspond to the edges of different optical images. Scale bar = 0.1 mm.

Phase contrast microscopy is preferable to bright field microscopy when observing the cell wall layer with high magnification (×600) as the specimen is thin, so the colour contrast is reduced (Abedini *et al*., 2015). Several images were acquired using a light microscope with a digital camera (DFC320, Leica Microsystems SAS, Nanterre, France) from the cambium to a distance of about 2 mm from it on the xylem side (*i.e*., with fully matured fibres), with a sufficient overlap to allow the image to be repositioned to accurately measure the distance of each cell from the cambium. The mean thickness of the S2 and G-layers was measured all along two radial rows using Matlab software (MathWorks Inc., Natick, Massachusetts, USA) according to the method of Yoshinaga *et al*. (2012). External contours of the lumen, S2 and G-layers were plotted by hand from images and their average thickness was calculated as (Abedini *et al*., 2015):

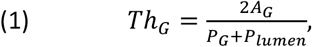

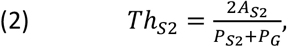

where *A_G_* and *P_G_* are the area and the external perimeter of G-layer, respectively, *A_S2_* and *P_S2_* are the area and the external perimeter of the S_2_ layer, respectively, and *P_lumen_* is the lumen perimeter. The data presented in the present article show the thickness of each layer normalized by the mean cell diameter, *D*, which was evaluated as 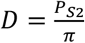. The advantage of working with relative thickness is that it allows the effect of the fibre ends on the cell wall thickness to be corrected (Okumura *et al*., 1977; Abedini *et al*., 2015).

### AFM PF-QNM measurements

Mechanical characterisation was performed with a Multimode 8 AFM (Bruker, Palaiseau, France) in PF-QNM imaging mode with a RTESPA-525-30 (Bruker) probe. The spring constant of the probe was calibrated by Bruker using a laser Doppler vibrometer with a value of 158 N/m. The initial tip radius, 33 nm (controlled by Bruker), was checked after adjusting the cantilever deflection sensitivity on sapphire and corrected to 40 nm to obtain the right range of indentation modulus on the centre of DuPont^™^ K48 Kevlar^®^ fibres (~20 GPa) embedded in Epofix (Struers SAS, Champigny sur Marne, France) epoxy resin (~4 GPa), as described in Arnould *et al*. (2017). The value of the tip radius was checked indirectly all along the measurements, and if necessary corrected using the above-mentioned calibration sample, by ensuring that the indentation modulus and the adhesion force in the embedding resin surrounding the wood sample and within the lumen in the cambial area remained constant. After all the measurements, the final tip radius was 120 nm. The applied maximum load was set at 200 nN for all the measurements, the vertical motion for force-distance curves was set at a frequency of 2 kHz, and the fast scan rate was such that the scan speed was always equal to 8 μm/s regardless of the size of the image (512 × 512 pixels), with a scan axis angle of 90°.

The force-distance curves obtained were automatically adjusted by a Derjaguin-Muller-Toporov (DMT) contact model (Derjaguin *et al*., 1975) to obtain the indentation modulus using Nanoscope Analysis software (Bruker, Palaiseau, France), with an assumed spherical tip, a flat sample surface, and taking the measured adhesion force into account. This model is one of the simplest and is suitable for vitreous polymer resin and all wood cell wall layers, considering the relatively low values of their Tabor parameter (Johnson and Greenwood, 1997; Xu *et al*., 2007). The discernible layers, *i.e*., layers that are thick enough to avoids the measurement being influenced by edge or topography effects, are the cell corner middle lamella (CCML), S_1_ with the primary wall (*i.e*., S_1_-P, as in most cases, these two layers are almost impossible to distinguish), S_2_ and G-layers. For each of these layers, the indentation modulus distribution was obtained using Gwyddion freeware (http://gwyddion.net/), see Fig. S3 in the Supplementary Material. This distribution can be adjusted with a Gaussian function that gives the value at the maximum of the distribution (*i.e*., mode or most frequent value in the dataset) and the standard deviation of the indentation modulus. Measurements were made on three different radial rows of developing cells in the wood sample, one after the other, always starting from the cambium and continuing up to a distance of about 1.7 mm away, with two overlapping sets of measurements for the first row to check the stability and repeatability of the measurements. Twenty-four different positions (and thus cells) were measured in the two first radial rows and 12 positions in the last row. As soon as it was visible, the thickness of the S_2_ and G layers was measured using the same protocol as for the optical images, Eqs. (1) and (2). To complete our study, and to have a reference, we measured the indentation modulus and the thickness of the cell wall layers in three normal wood cells (one per radial row) that had differentiated before the tree was tilted and were therefore devoid of a G-layer. All the data were assembled using Matlab software (The MathWorks Inc., Natick, Massachusetts, USA). Finally, the AFM values were checked by nanoindentation measurements on a few cells located 700 μm from the cambium using iNano KLA nanoindenter (Scientec, Les Ulis, France) in mapping mode (NanoBlitz) on a 200 × 200 μm (20 × 20 pixels) area, with a maximum force of 0.1 mN and a loading frequency of 1 Hz.

## Results

### Mapping the indentation modulus of developing fibres

The AFM measurements provided at least a map of the sample topography and a map of the indentation modulus. Examples of typical maps obtained for a cell are given in Fig. 2, at a distance of 740 μm from the cambium (first radial row). The different layers of the cell wall (cell corner middle lamella CCML, primary cell wall P, secondary cell wall S_1_, S_2_ and G-layers) are clearly identifiable on the indentation modulus map due to their different elastic mechanical properties. Note that part of the cell contents in the lumen are identifiable (Fig. 2b), while they are not visible in the topography (Fig. 2a). The different cell wall layers are also quite easy to distinguish on the topography map because of the slight change in height between each layer. The height is almost uniform within the G-layer, middle lamella and embedding resin in the lumen, whereas it varies around the circumference in the Si-P and S2-layers. These variations are the opposite in the S_1_-P and S_2_ (S_1_-P is high when S_2_ is low) and these extreme values were obtained perpendicular to the cutting direction (white dashed arrow in Fig. 2a). These observations are typical of a cutting effect as previously described in Arnould and Arinero (2015).

**Figure 2:**
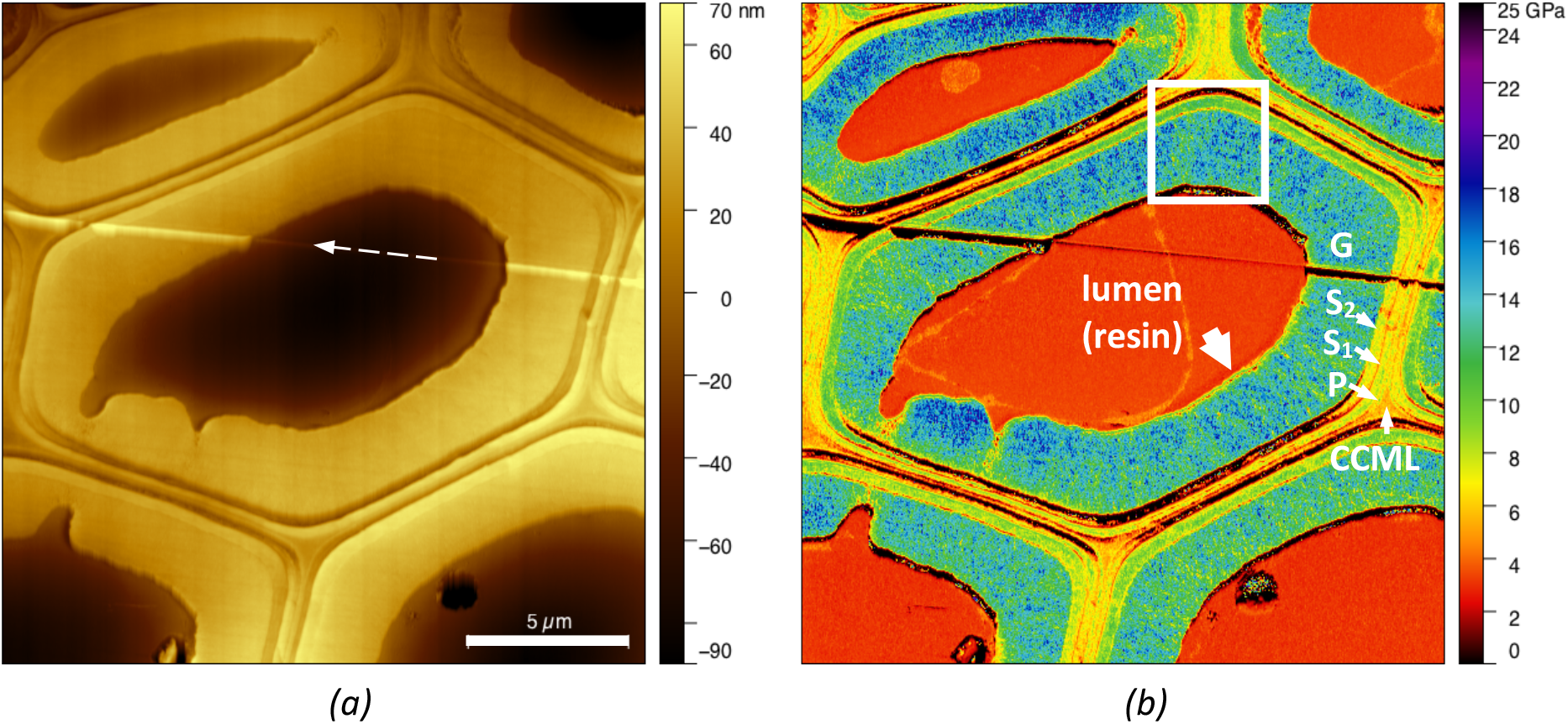
PF-QNM mapping of (a) topography and (b) indentation modulus of the cross section of a tension wood fibre at 740 μm from the cambium (first radial row). The different layers are identified: P stands for primary wall and CCML for cell corner middle lamella. The lumen of the cell was filled with LR-White resin. The white dashed arrow in (a) shows the microtome cutting direction (following a scratch line due to imperfections of the diamond knife), the thick white arrow in (b) points to a thin softer sub-layer that is more visible in Fig. 4, which is an enlargement of the white upper box in (b).

Moreover, we observed limited orthoradial variations in the indentation modulus of the S_2_-layer around the cells. This proves that the wood fibres are rather well oriented perpendicular to the cutting direction and that there will be little (or even no) bias in the interpretation of the measurements due to sample misalignment (Arnould and Arinero, 2015).

Fig. 3 shows the mechanical maps of all the cells measured in the first radial row. Progressive thickening of the cell wall results in the appearance of the different layers of the secondary wall: the first distinguishable S_2_ appears around 50 μm from the cambium (map with the green border in Fig. 3) and first distinguishable G-layer around 230 μm from the cambium (map with the blue border in Fig. 3).

**Figure 3:**
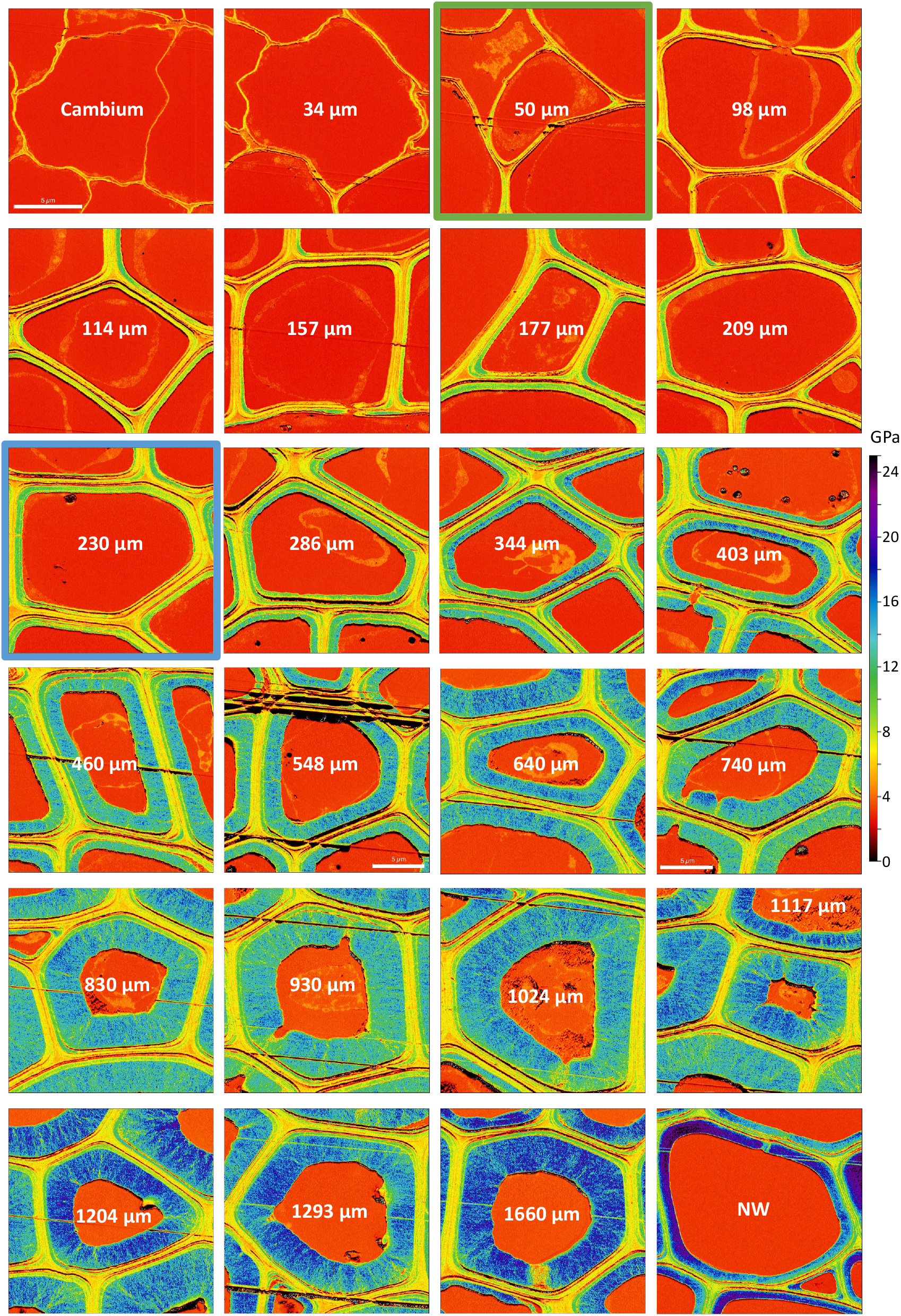
Indentation modulus maps of the different cells measured in the first radial row. The white number in the lumen refers to the distance of the cell from the cambium, the cells are arranged in rows from left to right and from top to bottom, with the cambium always on the left. The last map on the bottom right shows a normal wood (NW) cell, here before tilting (Fig. 1). The map at 50 μm (green border) is the first map with a distinguishable S2-layer. The map at 230 μm (blue border) is the first map with a distinguishable G-layer. Except for the maps at 548 and 740 μm, the size of the maps is same in all the images. Scale bar = 5 μm.

A continuous increase in the indentation modulus of the embedding resin is visible in the lumen from 2.7±0.1 GPa in the cambium to 3.4±0.2 GPa at 1.7 mm. This increase was not observed in the embedding resin outside the wood sample where the indentation modulus remained equal to around 2.7±0.1 GPa in all the measurements. Moreover, immediate measurement of the indentation modulus of the embedding resin in the lumen of cells in the cambium, taken just after the last measured cell in a given row, showed a return back to the initial value of 2.7±0.1 GPa.

The indentation modulus obtained for the S2-layer of normal wood cells 2 mm from the cambium, was around 16.9±5.5 GPa and its relative thickness was around 0.055 (see NW in Fig. 3). A more pronounced variation of the indentation modulus was observed in the S2-layer of this cell, which is probably due to a slight misorientation of the fibre with respect to the surface, as already described in Arnould and Arinero (2015).

The indentation moduli of the other layers were 7.5±1.2 for the CCML and 8.2±3.1 GPa for the S_1_-layer, while the indentation modulus in the embedding resin in the lumen was 2.99±0.21, a value close to that recorded in the cambium or in the resin surrouding the wood sample. The indentation modulus was confirmed by nanoindentation in the embedding resin in the lumen and in the G-layer of a few cells 700 μm from the cambium with a value of 3.5±0.15 GPa and 13.5±1.3 GPa, respectively (see Table 1 for comparison).

**Table 1:** Comparison of the value of the indentation modulus (in GPa) in the different layers of developing and mature wood fibres in our study and in the literature.

| Reference | LR-White resin (lumen) | ML (CC) | S<sub>1</sub> | S<sub>2</sub> | G |
| --- | --- | --- | --- | --- | --- |
| This study, developing tension wood (740 µm, Figs. 2 and S3) | 3.10±0.29 | 5.4±1.0 | 6.5±1.4 | 8.3±2.2 | 13.0±3.1 |
| This study, mature tension wood (1660 µm, Fig. 3) | 3.35±0.27 | 5.9±1.0 | 6.7±1.2 | 8.2±2.6 | 16.5±3.3 |
| This study, mature normal wood (NW, Fig. 3) | 2.99±0.21 | 7.5±1.2 | 8.2±3.1 | 16.9±5.5 | n.a. |
| Normand *et al.* (2021) (*Populus euramericana*, “Robusta” and “I214” blend) | 3.9±1.8 | 9.9±1.2 | 11.3±0.3 | 16.4±0.4 | 16.8±0.5 |
| Clair *et al.* (2003) (*Quercus ilex* L., no embedding) | n.a. | 5-7 | 8-9 | 9-10 | 10-12 |
| Arnould and Arinero (2015) (*Castanea sativa* Mill.) | 3.5±1.5 | 6±0.5 | n.a. | 13±0.5 | 15±1.5 |
| Liang *et al.* (2020) (poplar, no embedding) | n.a. | n.a. | 6.89-10.48 | 10.57-14.61 | 11.13-18.5 |
| Coste *et al.* (2021) (*Populus tremula* x *Populus alba*) | 4.5±0.9 | 10.7±2 | 16.0±3.8 | 18.2±3.5 | n.a. |

Overall stiffening of the G-layer with increased distance from the cambium was clearly visible. A radial pattern (radial lines in the cell wall) was also visible in the G-layer, as previously reported by Sell and Zimmermann (1998). Some ring lamellae were also visible within the cell wall layers *(e.g*., at 548, 740, 830, 930, 1024 and 1660 μm from the cambium in Fig. 3 and in the enlargement of Fig. 2b in Fig. S1 in the Supplementary Material). This last structural pattern is consistent with the radial layer-by-layer thickening of the wall and has been already reported, for example, in the S_2_-layer of wood fibres (Fahlén and Salmén, 2002; Casdorff *et al*., 2018), in the G-layer of most *Salicaceae* species except in the poplar genera (Ghislain *et al*.,2016), in mature (Hock, 1942) and developing G-layers of flax bast fibres (Arnould *et al*., 2017; Goudenhooft *et al*., 2018) and in mature hemp fibres with a G-layer (Coste *et al*., 2020).

At a distance from the cambium equal to or greater than 460 μm, a thin and soft sub-layer was visible on the lumen side at the border of the G-layer but only on the right side of the map (as shown in Fig. 2b). The fact that this sub-layer is only visible on the right side of all cells can be attributed to a cutting effect when the sample surface was prepared with the diamond knife, as the cutting direction is almost horizontal and proceeds from the right to the left (see Fig. 2a). As cutting effects are linked to the mechanical behaviour of the cell wall, this sub-layer reveals a different behaviour than the rest of the G-layer. The average indentation modulus of this sub-layer was around 8.2±2.6 GPa, close to the value of the early G-layer, at a distance of 230-286 μm from the cambium, and its thickness was around 100 nm in all cases. Fig. 4a gives a closer view of the G-layer at the top of the cell at 740 μm from the cambium (white box in Fig. 2b) and Fig. 4b is the adhesion map obtained by AFM. Although the sub-layer is not visible on the indentation map in Fig. 4a, a sub-layer with a thickness of around 100 nm and a lower adhesion force than the rest of the G-layer is also visible on the border of the lumen in Fig. 4b. We can assume that it is the same sub-layer as that observed on the right side of the indentation modulus maps. Moreover, its low adhesion force is close to that of the early G-layer (see Fig. S2 in the Supplementary Material).

**Figure 4:**
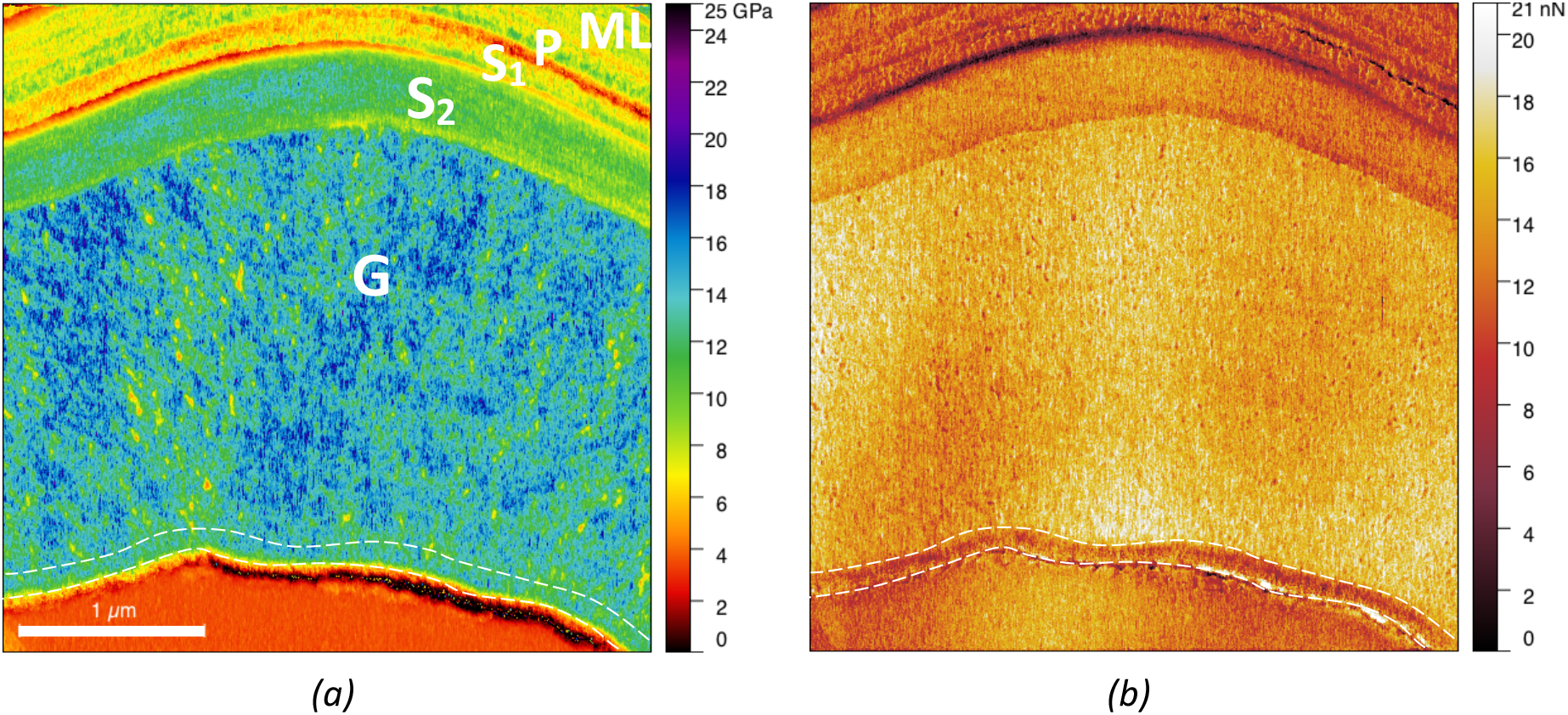
Close-up of the indentation map of a cell taken at a distance of 740 μm from the cambium corresponding to the white box in Fig. 2b, with the associated adhesion map (b), highlights a sub-G-layer with lower adhesion force close to the lumen.

To further investigate the kinetics of G-layer stiffening, we extracted six to ten radial profiles of the indentation modulus around the cell axis in the G-layer of six fibres situated at different distances from the cambium (Fig. 5). Each point in a radial profile is the average of the modulus over a width of 10 pixels. To reduce possible bias in the interpretation of the data caused by an edge effect due to cutting with the diamond knife or an effect of the area mechanically sensed by the tip (Sudharshan Phani and Oliver, 2019), we removed the first and last 100 nm from each profile (data points in grey in Fig. 5). In contrast to the indentation modulus map in Figs. 2b and 3, where no mechanical gradient is visible in the developing G-layers, here a gradient was always visible on the last 500 nm or so on the lumen side and became less pronounced with an increase in the distance from the cambium. The gradient completely disappears in the mature fibre (see Fig. 5 at 1 660 μm). It was not possible to determine whether such a gradient existed in the S2-layer because, even if it were present, it would be hidden by the effect of the apparent microfibril angle due to the slight misalignment of the sample (Arnould and Arinero, 2015).

**Figure 5:**
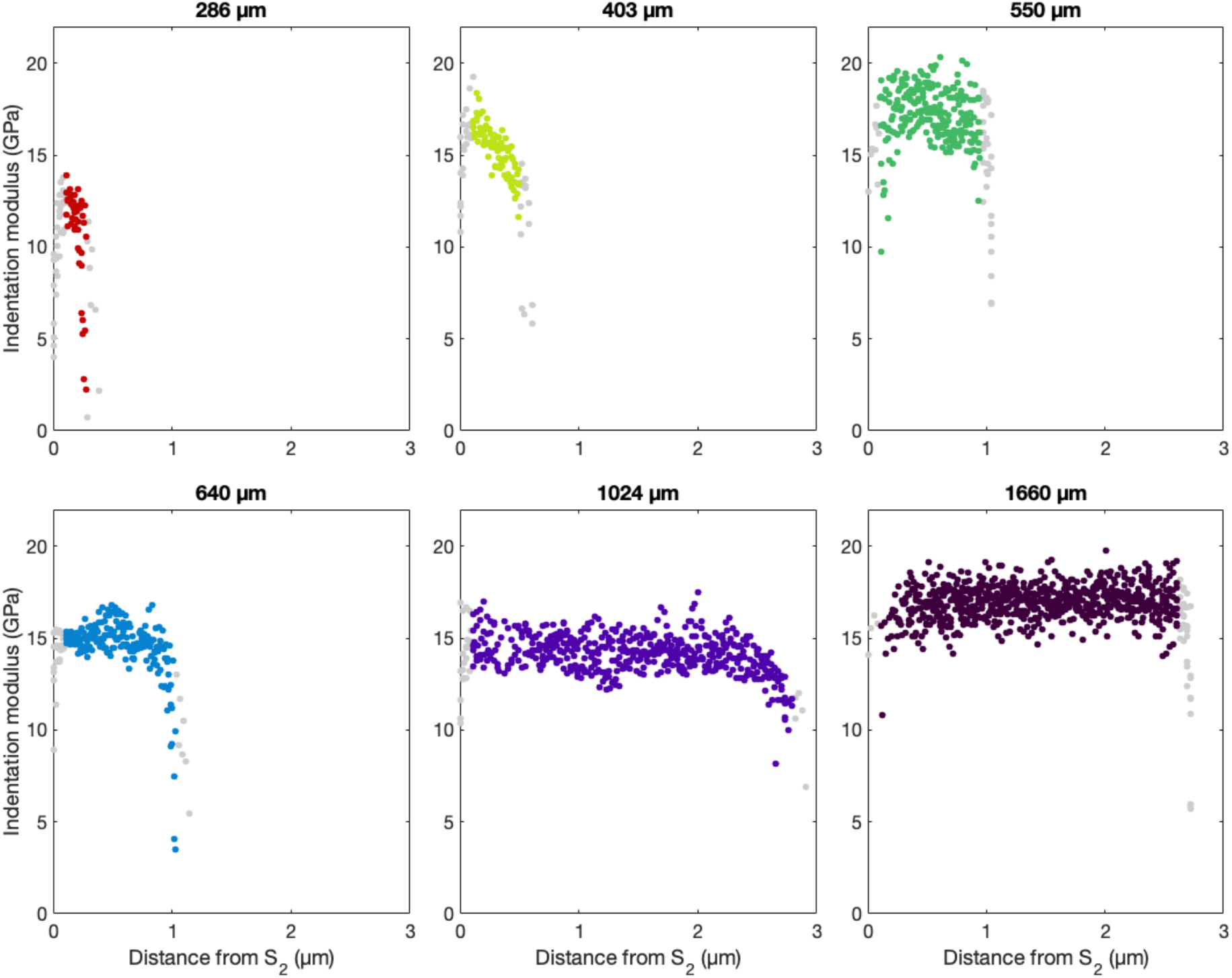
Observation of the occurrence of a radial mechanical gradient during the maturation of the G-layer obtained by extracting radial profiles all around the cell axis in this layer and plotting them as a function of the distance from the S_2_ layer, for fibres at six different distances from the cambium (value given at the top of each graph). The first and last 100 nm were removed from each profile (data points in grey) to avoid any bias due to possible measurement edge effects.

### Kinetics of global cell-wall layer thickening and stiffening

All the observations made above were also made in the 2^nd^ and 3^rd^ radial rows. Changes in the mode of the indentation modulus distribution in each layer (*e.g*., see the distribution of the indentation modulus in the different layers in Fig. 2b given in Fig. S3 in the Supplementary Material) are shown in Fig. 6, as a function of the distance from the cambium, together with the relative thickness of each layer. In Fig. 6, one point corresponds to one cell, whatever the radial rows, the continuous line corresponds to the mean trend adjusted on these points by a polynomial fit and the coloured ribbon to this fit shifted vertically by plus or minus the mean standard deviation on each layer of the cell wall.

**Figure 6:**
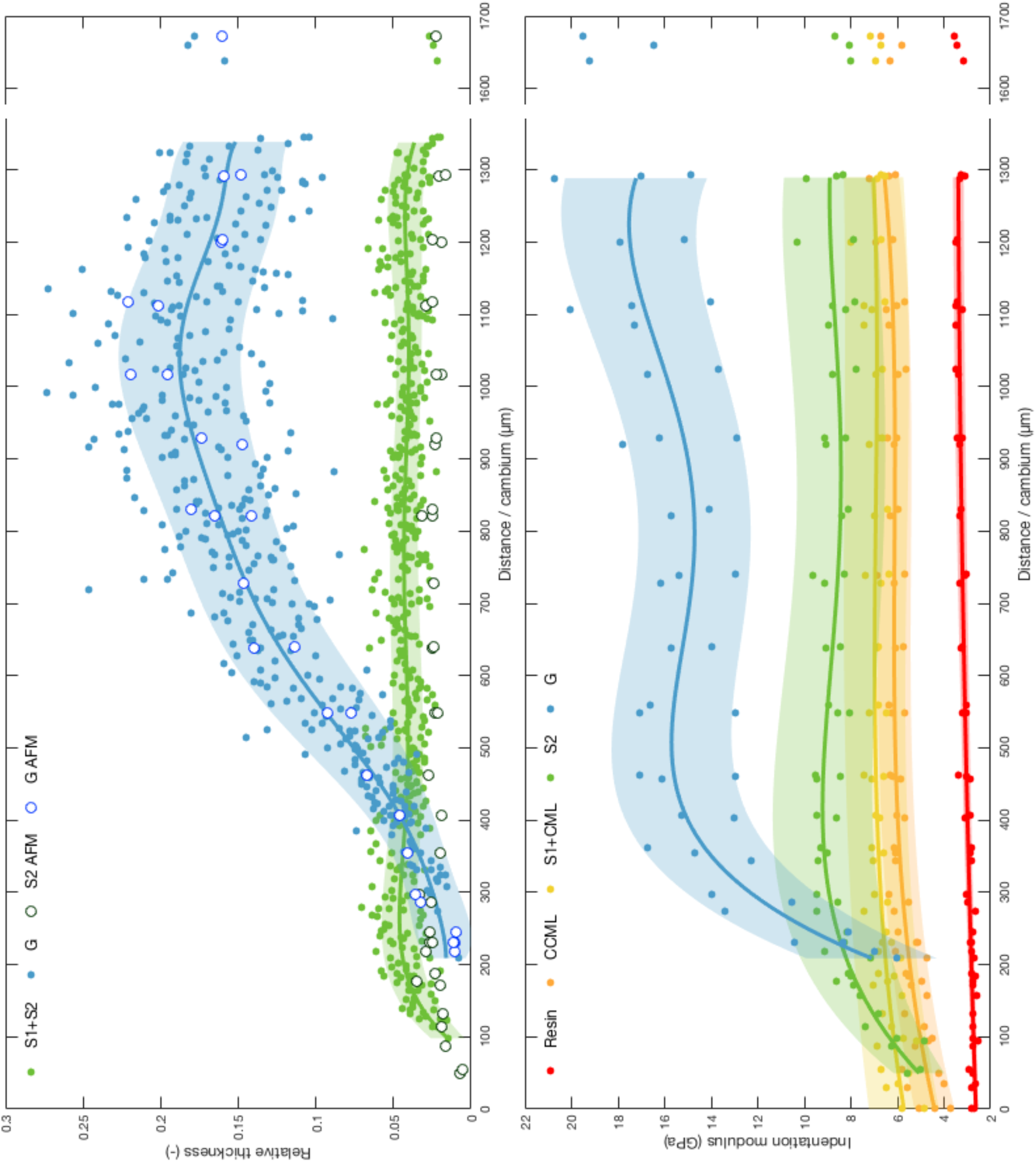
Variations in the relative thickness of the cell wall layers measured by optical microscopy (coloured dots) and AFM (empty circles) (top) and mode of the indentation modulus distribution (bottom), as a function of the distance from the cambium. The solid lines and the shaded areas show the mean tendency and standard deviation adjusted on these points.

In the case of the optical measurements of the thickness of the layers, it was not possible to separate the S_1_ and S_2_-layers, unlike for the AFM measurements. The measurements of relative thickness made by optical microscopy and AFM are consistent, but AFM enables detection of the appearance of the cell wall layer and its thickening earlier than optical microscopy. The thickness of the S_2_ alone obtained by AFM is thus logically smaller than S_1_+S_2_ obtained by light microscopy. The relative thickness of the S_2_-layer increases until around 200 μm from the cambium then decreases a little before reaching a stable value at a distance of around 500 μm from the cambium. The G-layers were first detected close to 200 μm from the cambium. The relative thickness of the G-layer increased linearly and stabilised near 1000 μm. Thus, the relative thickness of the S2-layer was slightly higher before the appearance of the G-layer.

A progressive increase in the indentation modulus of both the CCML (from 4.6±0.7 to 6.1±0.7 GPa) and the S_1_ layers (from 5.6±1.5 to 6.8±1.3 GPa) was observed until the end of the S_2_ stiffening, at around 350 μm from the cambium. The very first S_2_-layers had indentation moduli of 5.1±1.4 GPa and their stiffening and their thickening were initially synchronous. Later, when the S2-layers reached their final thickness, their indentation modulus continued to increase and finally reached a value of 8.7±2.0 GPa. All these layers continued to stiffen when the G-layers began to thicken. In contrast, the global stiffness of the G-layer reached an almost stable plateau (at around 500 μm from the cambium) long before it attained its final maximum thickness (at around 1000 μm from the cambium).

As already mentioned, as curves in Fig. 6 correspond to the mode of the indentation modulus distribution (*i.e*., the value at the maximum of the distribution or most frequent value, see Fig. S3 in the Supplementary Material), they do not reflect the gradient observed at about 500 nm from the edge of the G-layer on the lumen side due to the progressive maturation of a potentially freshly deposited sub-G-layer (Fig. 5). Furthermore, as shown in Fig. 5, the thickness of the G-layer at 550 μm from the cambium is such that most of the G-layer has completely stiffened, leading to the stabilised value of the indentation modulus reported in Fig. 6 for this distance from the cambium.

To compare the kinetics of the stiffening of the S_2_ and G-layers, Fig. 7 shows the normalized indentation modulus *(i.e*., the modulus from Fig. 6 divided by its mean maximum value) as a function of the distance from the cell where the layer concerned first appeared (*i.e*., 50 μm from the cambium for S_2_ and 230 μm for G-layers, Fig. 3). This figure shows that the kinetics of the two layers are quite similar, *i.e*., it took a distance of around 250 μm to globally reach their mature modulus. However, it appears to be faster for the G-layer as the change in modulus from the first deposited layer to the final mature one is larger.

**Figure 7:**
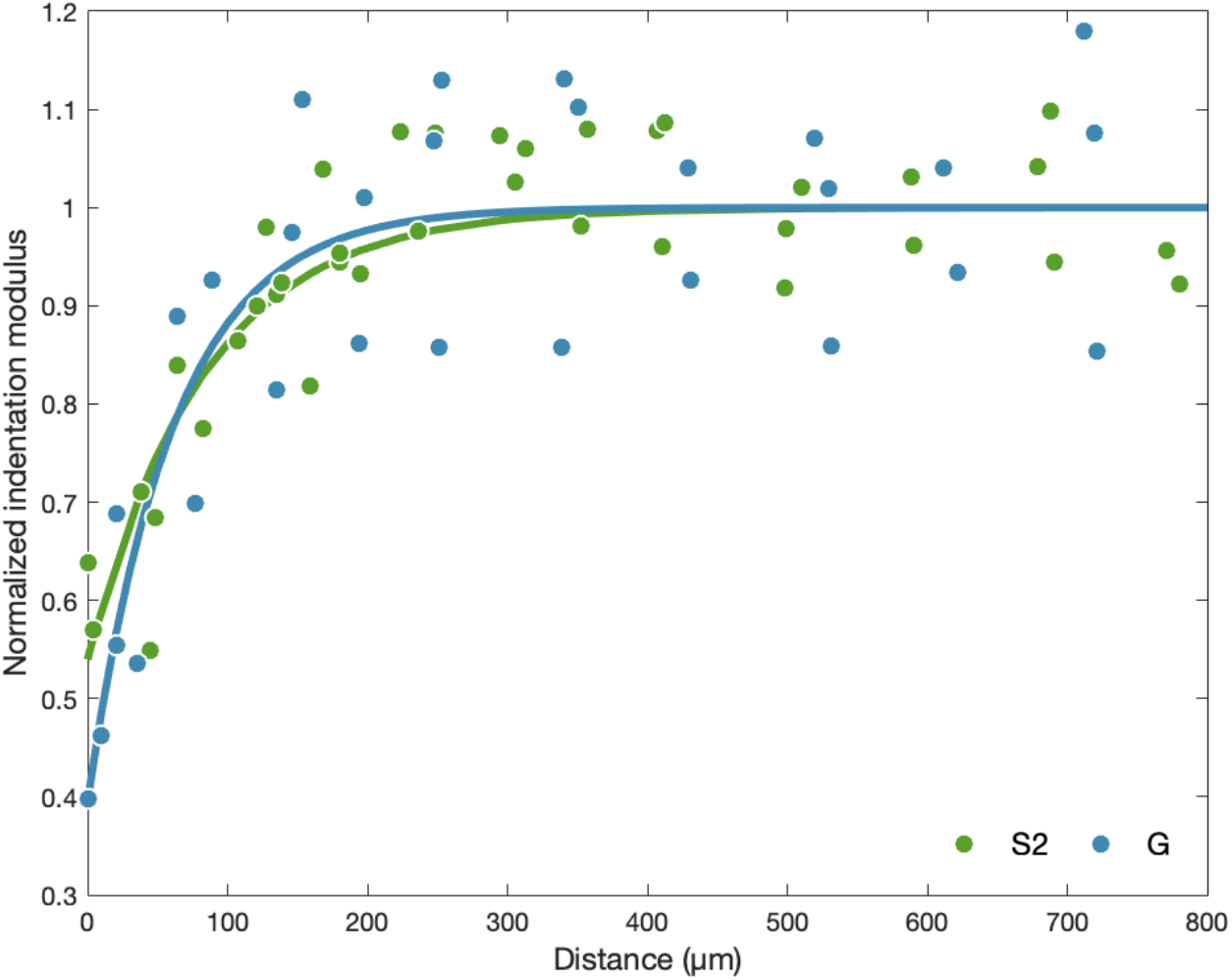
Normalized indentation modulus of the S2 and G-layers from Fig. 6 as a function of the distance from the cell where the layer concerned first appeared. The solid line corresponds to the mean value.

## Discussion

Our main results revealed: i) initial synchronous stiffening of the CML, S_1_ and S_2_-layers with the thickening of the S_2_-layers, which continues a little after the S_2_-layer has reached its final thickness when the G-layer starts to develop; ii) initial global stiffening of the G-layer synchronous with its thickening but stable global stiffness reached long before its final maximum thickness; iii) a stiffness gradient over of about 500 nm on the lumen side in the developing G-layer with a softer sub-layer at the lumen edge about 100 nm in thickness.

### Potential effects of sample preparation on the measurements

The different steps of sample preparation protocol made it impossible to keep the sample in its native *in planta* green state: we thus cannot rule out the possibility that modifications of the different layers of the cell wall during the ethanol exchange and resin embedding had some impacts on its mechanical properties but, for the reasons detailed below, we believe that we achieved a good compromise. Indeed, this preparation was necessary to ensure reliable mechanical measurements at small scale by AFM. Since all the measurements had to be comparable, this treatment minimised artifacts caused by roughness of the sample surface (Peaucelle, 2014). Indeed, mechanical measurements based on indentation require samples with a surface that is as flat as possible, compared to the radius of the AFM tip, to enable the use of reliable and simple contact mechanics models. These models are needed to extract the indentation modulus from the contact stiffness (Arnould and Arinero, 2015) or from the force-distance curves (Hermanowicz *et al*., 2014). In addition, the AFM tip is very brittle and surface roughness has to be as low as possible to reduce the risk of tip wear or breakage: this is especially important in the present study where we had to perform many measurements using the same probe to limit measurement bias or drift. Moreover, AFM measurements at such a small scale are only sensitive to the very near sample surface. Damage during preparation of the sample surface should therefore be reduced to the strict minimum. In addition, as we expected to find evidence for the existence of a mechanical gradient during the thickening of the cell wall layers, we had to begin taking measurements as close as possible to the cambium, where the cell wall is very thin and soft. This is only possible when the sample has been previously embedded to avoid, or at least reduce, deformation and damage during cutting and measurements. In addition, cell wall thickening progresses from the lumen side of the cell wall and, without embedding, measurements made close to the lumen would be highly modified due to border effects (Jakes *et al*., 2008; Jakes *et al*., 2009) unless the lumen is filled with a sufficiently stiff substance such as resin. Finally, these embedding steps reduce cell wall layer deformation during the cutting process and avoid swelling, detachment and collapse of the G-layer commonly observed after stress release (Clair *et al*., 2005a; 2005b).

Other studies have shown that LR-White embedding resin has little impact on the mechanical properties of the cell wall due to very limited penetration into the cell wall of normal wood (Coste *et al*., 2021) and *a priori* in the G-layers of tension wood (Arnould and Arinero, 2015) and of other similar fibre cell walls such as in flax (Arnould *et al*, 2017) and hemp (Coste *et al*., 2020). What is more, the use of ethanol is expected to cause only slight deformation of the wall. For example, Chang *et al*. (2012) showed that ethanol dehydration produced longitudinal macroscopic shrinkage of only 0.2% and volumetric swelling of only 0.5%. It is possible to avoid ethanol dehydration by drying the sample at moderate temperature just before embedding (Konnerth *et al*.,2008). However, in the present biomechanical context with the G-layer, such a drying step would lead to very significant changes in the cell wall ultrastructure, such as mesoporosity collapse (Clair *et al*., 2008).

The main impact of sample preparation on the mechanical properties of the cell wall is in fact its potential effects on the moisture content of the different layers. Indeed, sample preparation probably modified moisture content from a green state to close to an air-dry state. The effect of moisture content on the mechanical properties of the different cell wall layers has already been measured by nanoindentation in the cell corner middle lamella and the S2-layer of different woody species using samples that were embedded (Wagner *et al*., 2015) or not (Bertinetti *et al*., 2015; Meng *et al*., 2015). These studies revealed a similar trend with a reduction of the indentation modulus from one third to one half for the S2-layer and at least one half for CCML, between an air-dry and saturated state. A more recent study (Coste *et al*., 2020), using AFM PF-QNM in similar conditions to those used in our study, focused on the effect of the moisture content on the mechanical properties of hemp sclerenchyma fibres (containing a thick G-layer with similar characteristics to those of the tension wood G-layer) and xylem fibres. In their study, AFM measurements of all the cell wall layers revealed no major differences between layers, with a reduction of the indentation modulus of about one half when the relative humidity varied from 13% to 83%. If we extrapolate these variations to our study, the indentation modulus values reported here are overestimated compared to the values *in planta* but the relative differences observed between layers, or within a layer (gradient), are most probably comparable to what happens in the tree.

### Indentation modulus and its variations in the different layers of the cell wall

We observed an increase in the indentation modulus of the embedding resin in the lumen, with increasing distance from the cambium, but it returns back to values measured in the cambial zone in the normal wood (before tilting) cells lumen. The origin of this increase during fibre maturation is not yet understood but is unlikely to be due to wear of the AFM tip as demonstrated by the repeatability of the measurements in the cambial cells performed after measurements of each row, which were also identical to those obtained at the end of all measurements in the lumen of the normal wood cells or in the resin outside the sample. Stiffening thus appears to be associated with the change in the contents of the lumen with the maturation of the fibres (as shown in Fig. 3). In cambial cells, the plasma membrane and cytoplasm are bound to the inner part of the cell wall. Cambial cells are highly vacuolated, and the large vacuole pushes the cell organelles outwards. There is therefore little material inside the lumen (vacuole contents), which may explain why the indentation modulus measured in the resin in the centre of cambial cells is close to that measured in normal wood cells that have lost all their cell contents. Finally, Table 1 shows that our LR-White indentation modulus values were the lowest when compared to other authors’ data, but were confirmed by nanoindentation using iNano KLA nanoindenter. This is probably due to differences in the calibration procedure between laboratories or to the variability of the resin itself, as different grades (soft, medium, and hard) of this resin are available.

The values of the indentation modulus in the different layers and the embedding resin are consistent with the (rather scattered) AFM data or nanoindentation measurements of wood cell walls available in the literature (Arnould and Arinero, 2015; Clair *et al*., 2003; Coste *et al*., 2021; Eder *et al*., 2013; Liang *et al*., 2020; Normand *et al*., 2021), although in the low range compared to data in the literature on the G-layer of poplar or tension wood (see Table 1). These low values can be partly explained by the young age of the tree used in our study (less than 3-month old). Indeed, the juvenile wood is known for its high microfibril angle (MFA) in the S2-layer and its low cellulose content (Luo *et al*., 2021). The values of the indentation modulus in the G-layer of a mature cell increased to around 18.3±3.1 GPa on average (see Fig. 6), a value in the same range of the ones cited in the literature (Table 1).

The low value obtained for the mature S_2_-layer in the tension wood area compared to the value in normal wood can be explained by a marked difference in MFA between the S_2_-layers of normal wood (with a low MFA and therefore a high indentation modulus) and the S_2_-layers of tension wood (with a high MFA and therefore a small indentation modulus, Eder *et al*., 2013; Jäger *et al*., 2011). To explain this difference (equal to a factor of about 2) between the indentation moduli, we can roughly estimate from published data that the MFA is around 5-10° in normal wood whereas it is 30-40° in the S_2_ of tension wood (Arnould and Arinero, 2015; Jäger *et al*., 2011). This is also in agreement with the value of MFA reported for the S2-layer in tension wood for poplar by Goswami et al. (2008). Thus, using the data on the effect of MFA on the indentation modulus in Figure 3 of Jäger et al. (2011), a change from 10° to 40° of MFA results in a reduction in the indentation modulus of about 40%. Likewise, the order of magnitude of the values of indentation modulus obtained for the different layers of normal wood is in agreement with other literature data (Table 1).

### Kinetics of global thickening and stiffening of the cell-wall layers

The CCML, S_1_ and S_2_-layers continued to stiffen while the G-layer was developing (Fig. 6). This is in agreement with the fact that the lignification of S_1_, S_2_-layers and CCML occurs during the formation of the G-layer (Yoshinaga *et al*., 2012). This lignification after the G-layer starts to thicken may be explained by the presence of additional matrix material that has been transported through the existing wall. Alternatively, some precursors may already be present and are used in biochemical reactions that continue during the deposition of the G-layer. The effect of lignification on the mechanical properties of the cell wall is not yet well understood, with different studies sometimes reporting conflicting results, but recent studies tend to confirm the hypothesis that lignification mainly affects the shear modulus and the strength of the matrix (Özparpucu *et al*., 2017; 2019), with higher content leading to a higher modulus and greater strength. The indentation modulus is sensitive to the longitudinal modulus but also to the transverse and shear moduli (Jäger *et al*.,2011), which are mainly influenced by the cell wall matrix. Therefore, when lignification modifies the cell wall matrix properties, this results in a significant change in the indentation modulus, as already shown by nanoindentation (Gindl *et al*., 2002). Finally, Fig. 7 shows that the stiffening kinetics appear similar although faster in the G-layer than in the S_2_-layers suggesting that the physical and chemical changes or reactions at work during cell wall maturation are faster in the G-layer *(e.g*., microfibrils aggregation or gelatinous matrix swelling (Alméras and Clair, 2016)) than in the S_2_-layer (*e.g*., lignification).

The fact that the relative thickness of the S_2_-layer decreases slightly when the G-layer is starting to develop has already been observed. For example, Abedini et al. (2015) reported that this is a common trend throughout the growing season in both normal and tension wood of poplar trees. Moreover, the changes and mature value of the relative thickness of the G and S2 layers in Abedini *et al*. (2015), Chang *et al*. (2015) and Clair *et al*. (2011) are similar to our measurements. We therefore assume that we can use the relative thickening of the different wall layer as a common spatial reference to link different studies. If we combine our results with those of previous studies, the G-layer appears to synchronously stabilise its thickness, whole indentation modulus (*i.e*., no more radial gradient), meso-pore size (Chang *et al*., 2015) and cellulose tensile strain (Clair *et al*., 2011) at the end of the maturation. These observations suggest that the different changes involved in the maturation process of the G-layer start, evolve and end at approximately the same fibre development stage. These physico-chemical observations now need to be coupled with biochemical analyses to better understand the mechanisms involved in G-layer maturation, and possibly to establish relationships between matrix stiffening, bridging between microfibrils and wall compaction (Alméras and Clair, 2016; Gorshkova *et al*., 2015; Mellerowicz and Gorshkova, 2012).

According to the radial profiles of the indentation modulus (Fig. 5), a smooth mechanical gradient occurs in immature G-layer at less than 0.5 μm on the lumen side with a small sublayer of about 100 nm. This sublayer appears to be as dense as the mature part of the layer and could be either a freshly deposited immature G-layer or part of the periplasmic area still bound to the layer. Indeed, periplasmic area, located between the inner part of the G-layer and the plasma membrane, is the scene of intense biochemical processes, see Fig. 2 in Pilate *et al*. (2004), Fig. 5 in Guedes *et al*. (2017) or Fig. 7 in Decou *et al*. (2020). In contrast, flax bast fibres exhibit a strong mechanical gradient with a thick immature, loose and soft G-layer, called G_n_ (Gorshkova and Morvan, 2006; Gorshkova *et al*., 2010). Evidence for the presence of this thick G_n_-layer has also been provided in flax xylem tension wood fibres (Petrova *et al*., 2021). Interestingly, the indentation modulus is similar, or even a little bit higher, in flax G-layers than in mature poplar G-layers, while the average indentation modulus is in the same range in flax G_n_-layers, in immature poplar G-layers in fibres close to the cambium and in inner sub-layers observed in more developed G-fibres.

In a typical developing flax fibre, both indentation modulus (Arnould *et al*., 2017; Goudenhooft *et al*., 2018) and adhesion force exhibit a sharp transition between G and G_n_-layers as shown in Fig. 8. However, the sublayers observed as lamellae in the Gn have indentation modulus and adhesion force similar to those measured in the G-layer. These lamellae are separated by bands whose indentation modulus is close to that of the resin, but with a lower adhesion force. This lamellar arrangement is not observed in poplar, even though ring lamellae structure of this type is sometimes discernible in the mature part of the G-layer *(e.g*., see cells at a distance of 548, 740, 830, 930, 1 024 and 1 660 μm from the cambium in Figs. 3 and S1 in the Supplementary Material). The most significant structure in the poplar G-layer appears as radial bands (*e.g*., see tension wood fibres at a distance of more than 740 μm in Fig. 3). This pattern may reflect biological organisation, but we cannot exclude a possible consequence of a slight shrinkage of the G-layer during dehydration with ethanol (Fang *et al*., 2007).

**Figure 8:**
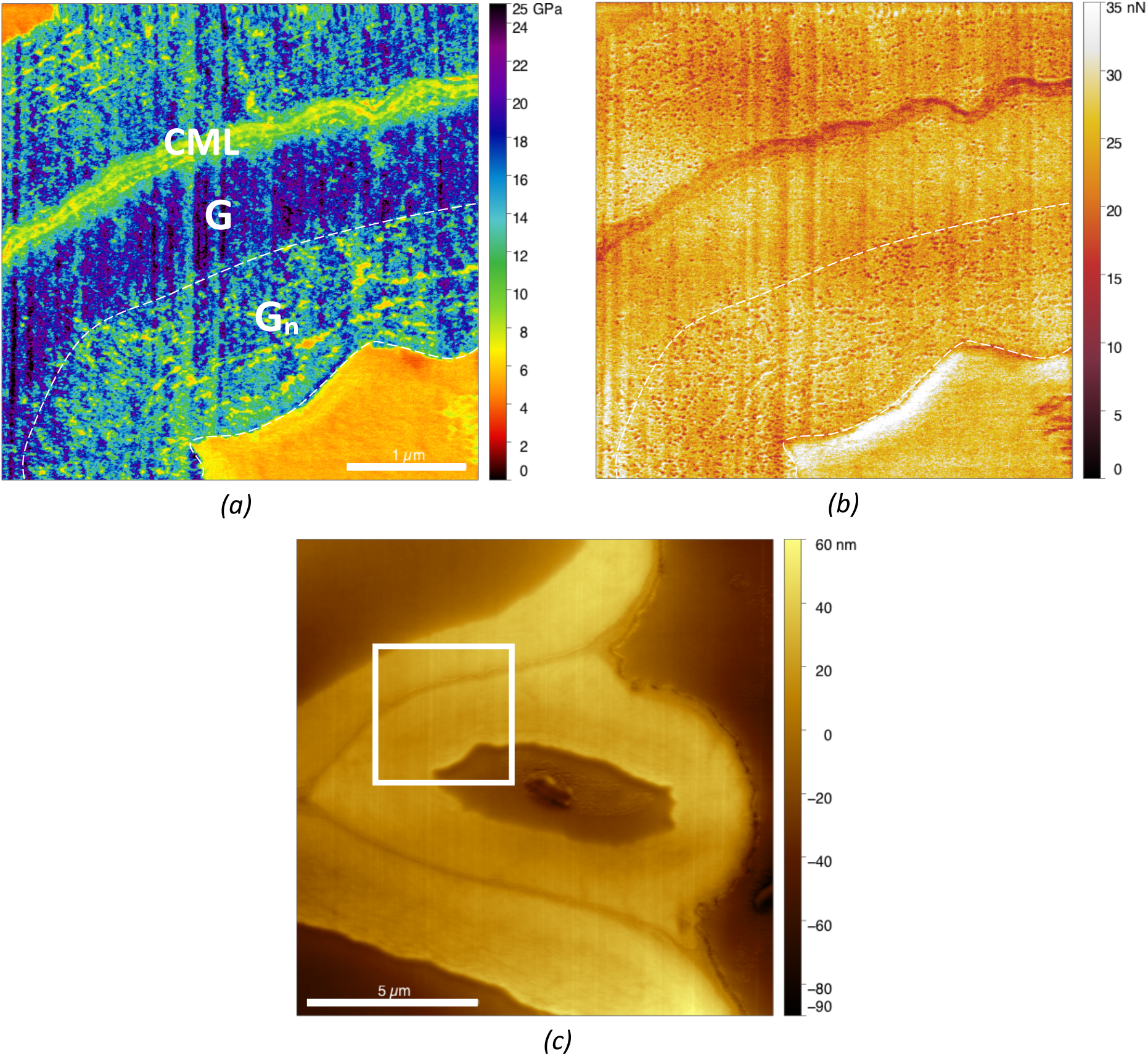
Comparison of the G and G_n_-layers in developing flax bast fibre (60 days, half height of the stem) adapted from Arnould *et al*. (2017): a) indentation modulus map and b) adhesion map corresponding to the white box in the topography image (c).

Note that it is not possible to compare the absolute value of adhesion forces obtained in the present study (Fig. 4b) with the values obtained in Arnould *et al*. (2017) (in Fig. 8b) as this force depends to a great extent on the shape of the tip and on the surface roughness of the material, which were not the same (see for example the difference in adhesion forces of the embedding resin in the lumen in the two studies, even though the same resin was used).

Although the G-layer of tension wood and the G-layer of flax are biochemically, ultrastructurally and mechanically similar (Coste *et al*., 2020; Guedes *et al*., 2017; Gorshkova and Morvan, 2006; Gorshkova *et al*.,2018; Petrova *et al*., 2021), they clearly differ in the kinetic of their development and maturation, as summarised in Fig. 9. Indeed, in flax, a thick and loose multilayered G_n_-layer stiffens and densifies abruptly, whereas, in poplar, it is a thin and dense immature layer that stiffens gradually. Further complementary analyses including immunochemistry need to be done to clarify the origin of these differences.

**Figure 9:**
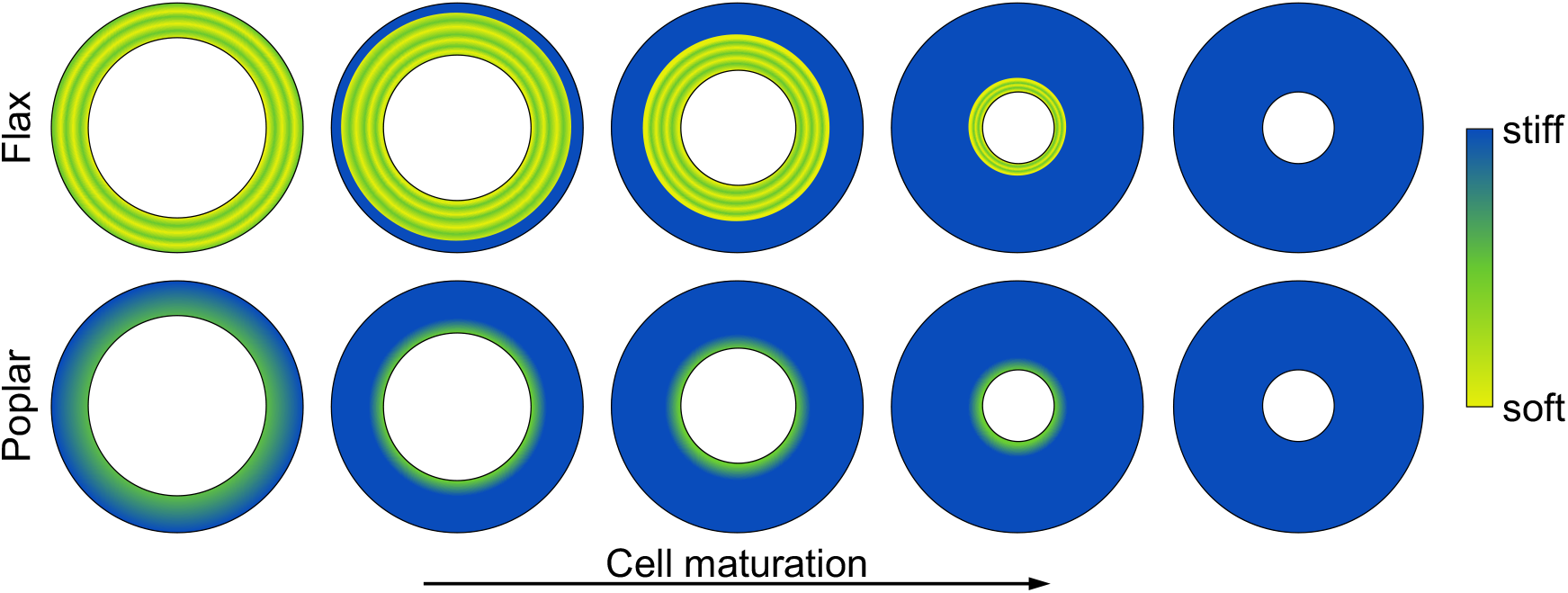
Comparative scheme of the maturation (thickening and stiffening) of the G-layer of flax and poplar.

## Conclusion

The use of AFM makes it possible to measure simultaneously the stiffening and thickening kinetics of different cell wall layers: this provides novel and precious insight into the kinetics of the maturation of any kinds of wood fibre. In this study, we applied this technique to poplar tension wood fibres containing a G-layer: this revealed that the G-layer reaches its near final stiffness long before its final thickness. In addition, we found evidenc of a radial mechanical gradient localised at the lumen periphery that remains throughout the thickening and disappears very late in mature G-layers. This contrasts with the maturation kinetics of the other cell wall layers, where thickening and stiffening are mostly synchronous. Finally, although the G-layer in poplar tension wood fibres and in flax phloem fibres are biochemically, ultrastructurally and mechanically similar, it is clear that they differ in the kinetic of their development and maturation.

The data collected in this study is not sufficient on its own to discriminate among the hypothetical mechanisms of maturation stress generation, reviewed in Alméras and Clair (2016). In this last article, the authors found that four mechanisms were admissible to explain stress generation in tension wood: (i) stress generation in amorphous cellulose domain in series with crystalline domain in the microfibrils, (ii) active binding of microfibrils by a (still unspecified) material, (iii) entrapment of material during microfibrils aggregation and cell wall compaction as suggested for flax bast fibres (Goudenhooft et al., 2018) (see Fig. 8b-c too) and (iv) swelling of the matrix in a connected cellulose network. In order to discriminate these different mechanisms, it is necessary to estimate their respective effects on the mechanical properties of the cell wall, and to estimate the resulting effect on the indentation modulus. Indentation modulus is a complex combination of different elastic parameters, particularly longitudinal, transverse and shear elastic properties (Jäger *et al*., 2011), the last two are particularly sensitive to the “matrix” moduli (*i.e*., matrix and binding between microfibrils). Thus, the first mechanism of maturation stress generation would probably have almost no effect on the cell wall mechanical properties, if not accompanied by a change in the matrix mechanical properties. Active binding and cellulose aggregation may have similar stiffening effect, in that it would lead to an increase in the shear and transverse elastic properties of the cell wall. Matrix swelling could lead to an apparent stiffer matrix (isotropic) property, but probably with a lighter effect than the two previous mechanisms. So, information about the change along the maturation process in the cell wall longitudinal, shear and transverse properties ratio is critical. Finally, more than one mechanism could be involved together or at different stage of the maturation. For example, it is possible that the slight and homogeneous increase in the indentation modulus that can be seen in Fig. 5 between 1024 μm and 1660 μm from the cambium and in Fig. 6 for a distance from the cambium greater than 900 μm, after the stiffening process described in the present study, was due to another stiffening mechanism. Further studies on the composition and structure of the G-layer (including, for example, immunochemistry) definitely need to be done in order to advance our knowledge.

## Abbreviations

AFM: Atomic force microscopy
CML: Compound Middle Lamella
CCML: Cell Corner Middle Lamella
MFA: Microfibril angle
PF-QNM: Peak-force quantitative nano-mechanics

## Acknowledgements

The authors are grateful to C. Assor (UMR IATE, Sup’Agro, INRAE Montpellier, France) for fruitful discussions and to D. Pellerin (ScienTec) for nanoindentation measurements. Version 4 of this preprint has been peer-reviewed and recommended by Peer Community In Forest and Wood Sciences (https://doi.org/10.24072/pci.forestwoodsci.100007).

## Data availability

The datasets used during the current study are freely available on the open repository website Zenodo: https://doi.org/10.5281/zenodo.6487575.

## Supplementary material

Supplementary material are available online: https://doi.org/10.1101/2021.09.23.461481.

