## Supplementary Figures for "Mechanical characterisation of the developing cell wall layers of tension wood fibres by Atomic Force Microscopy"

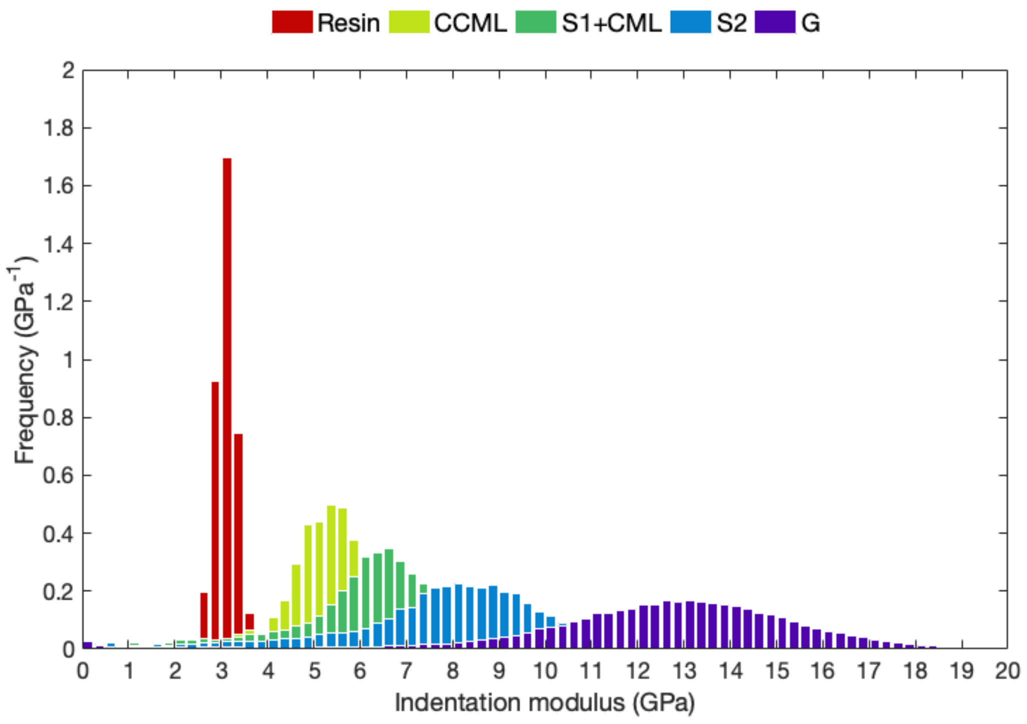


*Fig. S1. Distribution of the indentation modulus in the different layers of the tension wood fibre cell wall at 740 µm from the cambium (Fig. 2b). Resin corresponds to the embedding resin in the lumen.*

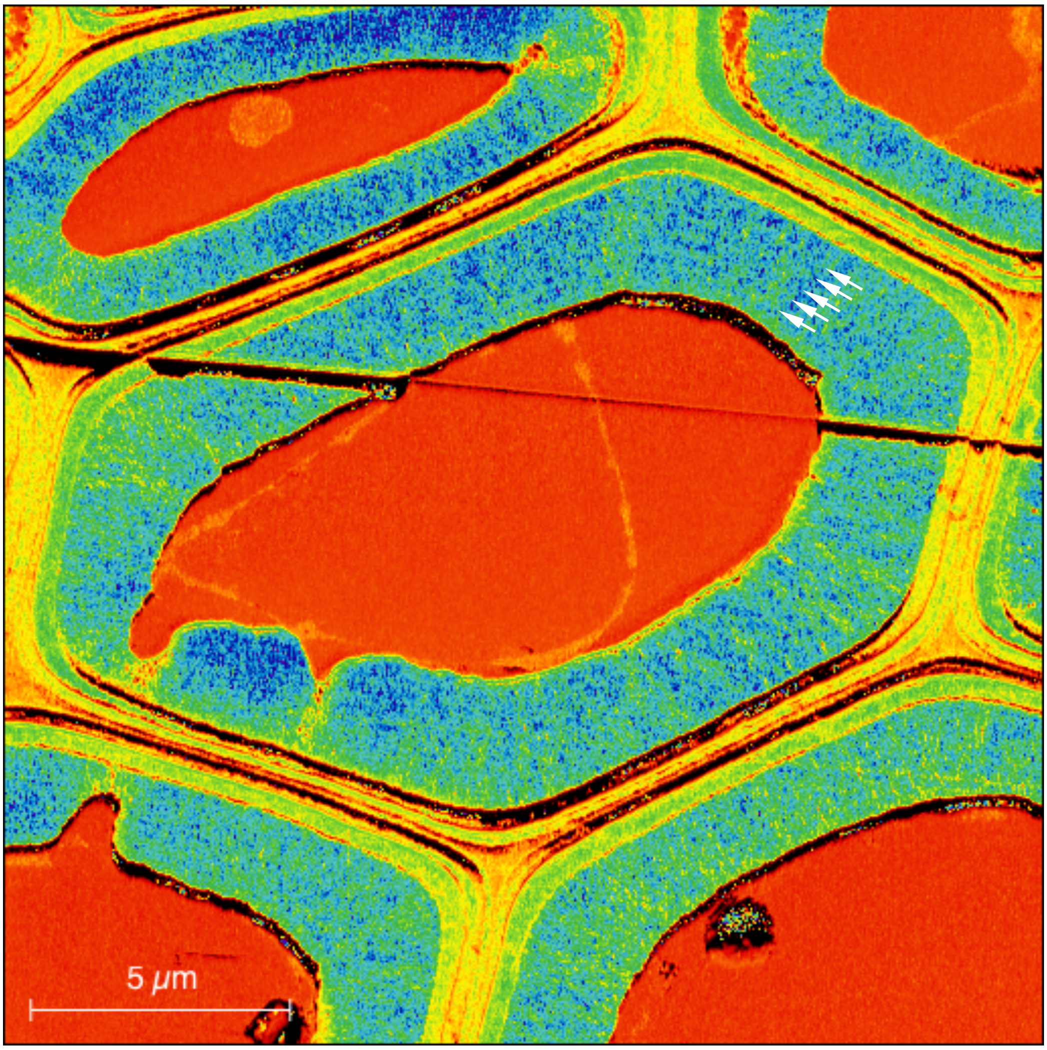


*Fig. S2. Enlargement of Fig. 2b to highlight the annular lamellar structure of the G-layer (white arrows) also visible at 548, 830, 930, 1 024 and 1 660 μm from the cambium in Fig. 3.*

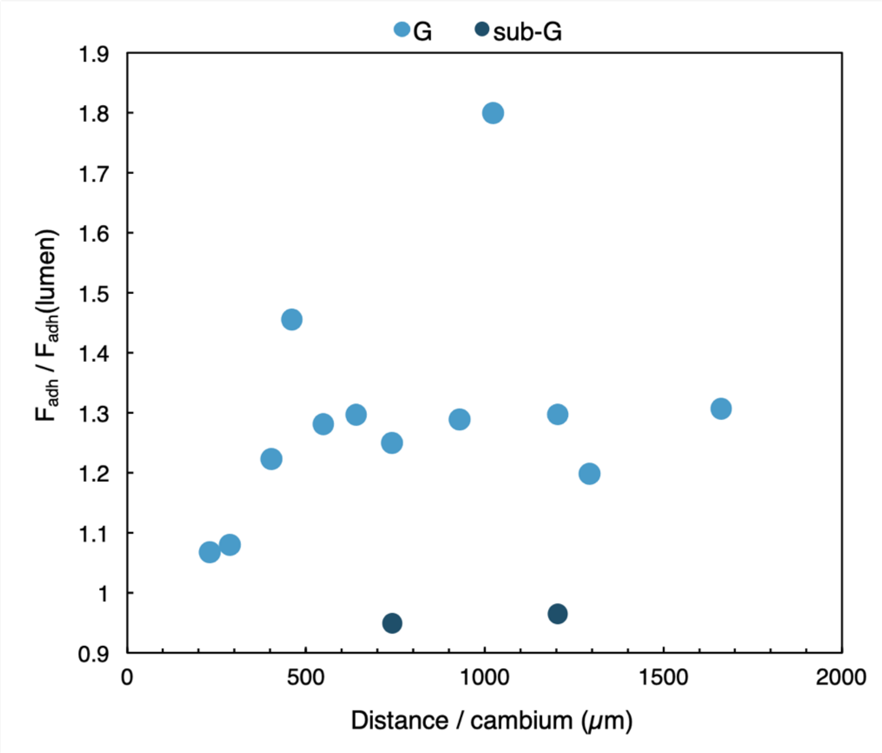


*Fig. S3. Changes in the average adhesion force in the G-layer (light blue) and sub-G-layer (dark blue) relative to the adhesion force on the embedding resin in the lumen.*
